# Development of a signal-integrating reporter to monitor mitochondria-ER contacts

**DOI:** 10.1101/2024.02.14.580376

**Authors:** Zheng Yang, David C. Chan

## Abstract

Mitochondria-ER contact sites (MERCS) serve as hotspots for important cellular processes, including calcium homeostasis, phospholipid homeostasis, mitochondria dynamics, and mitochondrial quality control. MERCS reporters based on complementation of GFP fragments have been designed to visualize MERCS in real-time, but we find that they do not accurately respond to changes in MERCS content. Here, we utilize split LacZ complementing fragments to develop the first MERCS reporter system (termed SpLacZ-MERCS) that continuously integrates the MERCS information within a cell and generates a fluorescent output. Our system exhibits good organelle targeting, no artifactual tethering, and effective, dynamic tracking of the MERCS level in single cells. The SpLacZ-MERCS reporter was validated by drug treatments and genetic perturbations known to affect mitochondria-ER contacts. The signal-integrating nature of SpLacZ-MERCS may enable systematic identification of genes and drugs that regulate mitochondria-ER interactions. Our successful application of the split LacZ complementation strategy to study MERCS may be extended to study other forms of inter-organellar crosstalk.

## Introduction

Mitochondria have major roles in promoting bioenergetic pathways, cell signaling, calcium homeostasis, and apoptosis^1–3^. In addition to true mitochondrial diseases, dysfunctional mitochondria have been linked to neurodegenerative diseases, including Parkinson’s disease, amyotrophic lateral sclerosis, and Huntington’s disease^2–5^. In recent years, inter-organellar crosstalk has emerged as a factor influencing the cellular roles of mitochondria^6–8^. In particular, the endoplasmic reticulum (ER) closely interacts with mitochondria and modulates cellular physiology. Such interactions occur at mitochondria–ER contact sites (MERCS), which are close appositions of mitochondria and ER with a distance of ∼10-50 nm^9^. These contacts regulate a number of cellular processes, including calcium homeostasis, lipid homeostasis, mitochondrial dynamics, and mitochondrial quality control^10–13^. Although some mitochondria-ER tethers have been identified^10,11,13–15^, much remains to be understood about the regulation of MERCS dynamics.

Given the physiological importance of mitochondria-ER interactions, it is critical to develop new tools to understand their molecular basis and cellular functions. Several studies have described reporters designed to identify MERCS in single cells^16,17^. Most current reporter systems employ biomolecular fluorescence complementation (BiFC). Based on the splitting of fluorescent proteins into two complementary fragments, these assays target the fragments individually to mitochondria and ER. In locations where the two organelles are in close proximity, the two protein fragments are reconstituted into a functional protein whose chromophore matures. Split-GFP, split-RFP, and split-Venus have been engineered to directly visualize the contact sites^18,19^. However, reconstituted fluorescent proteins form thermodynamically stable complexes^20^, and the long half-life for dissociation may perturb normal MERCS dynamics or cause artificial tethering of membranes. Double-dimerizing green fluorescent protein (ddGFP) has been used in a MERCS reporter^21^ but the signal-to-noise ratio is usually low in such systems. Fluorescence resonance energy transfer (FRET) and bioluminescence resonance energy transfer (BRET) have also been used to construct organellar contact sensors^16^. Because FRET/BRET methods do not require physical contact between the sensor partners, they avoid the potential problem of artificial tethering. However, these are proximity-based strategies and do not ensure that the signal arises from true physical contacts^22^.

All these reporters were designed to measure contacts sites at a specific point in time. However, MERCS are dynamic structures that assemble and disassemble depending on the physiological setting, and several cellular activities--organelle motility, organelle shaping, cell cycle--either regulate or depend on the dynamic nature of MERCS. A single time point measurement may not be an accurate representative of the overall MERCS content of a specific cell. As an example, mitochondrial fusion and fission events, which are regulated by MERCS, show stereotypical fluctuations as cells progress through the cell cycle^23^. Moreover, the essentially irreversible nature of GFP complementation raises the concern that BiFC-based reporters do not accurately respond to dynamic changes in the MERCS content. It would therefore be advantageous to have a MERCS reporter with integrative properties so that cells with overall high or low MERCS content can be accurately distinguished. Here, we designed the first MERCS reporter system with a fluorescent output that integrates information about the MERCS level over time. This reporter, termed SpLacZ-MERCS, utilizes α acceptor and α donor LacZ fragments targeted to the mitochondria and ER, respectively. The reporter accurately reads out the overall MERCS level and is responsive to dynamic fluctuations in mitochondria-ER interactions. We validated the ability of SpLacZ-MERCS to detect pharmacological and genetic perturbations known to affect MERCS. This reporter provides a new opportunity to investigate MERCS in high-throughput settings, including genome-wide perturbations and drug screening assays.

## Results and Discussion

### Identification of optimal LacZ fragments for MERCS reporter

In designing a new reporter to measure MERCS, we took advantage of the ability of weakly interacting fragments of β-galactosidase to reconstitute enzymatic activity. The bacterial LacZ gene, which encodes the enzyme β-galactosidase, exhibits α complementation, in which β-galactosidase containing an N-terminal truncation (termed α acceptor) can be complemented by an N-terminal peptide (termed α donor or α peptide) provided *in trans*^24^. Previous reports have described using α complementation with weakly interacting pairs of α donors and α acceptors to monitor protein-protein interactions in mammalian cells without artifactually enhancing their physical interaction^25–27^, a concern with split GFP approaches due to the high stability of the reconstituted complex. To adapt this system to study mitochondria-ER interactions in cultured cells, we started with a version of LacZ optimized for expression in mammalian cells^28^. Two types of α acceptors (LacZ Δα 6-36, LacZ Δα 6-78) were targeted to the mitochondrial outer membrane by fusion with the TOMM70 targeting sequence, and three types of α donor (LacZ α 1-75; LacZ α 1-141; LacZ α 1-782) were targeted to the ER membrane by fusion with the UBE2J2 targeting sequence. Glycine-serine linkers were included to provide polypeptide chain flexibility for refolding and enzymatic complementation (Figure 1A). Figure 1B illustrates the premise that these membrane-anchored split-LacZ fragments can form a complex and reconstitute β-galactosidase enzymatic activity only when the membranes of the mitochondria and ER are in close apposition. Due to the weak interaction of LacZ fragments, the protein complex could dissociate when a contact site disassembles.

**Figure 1.**
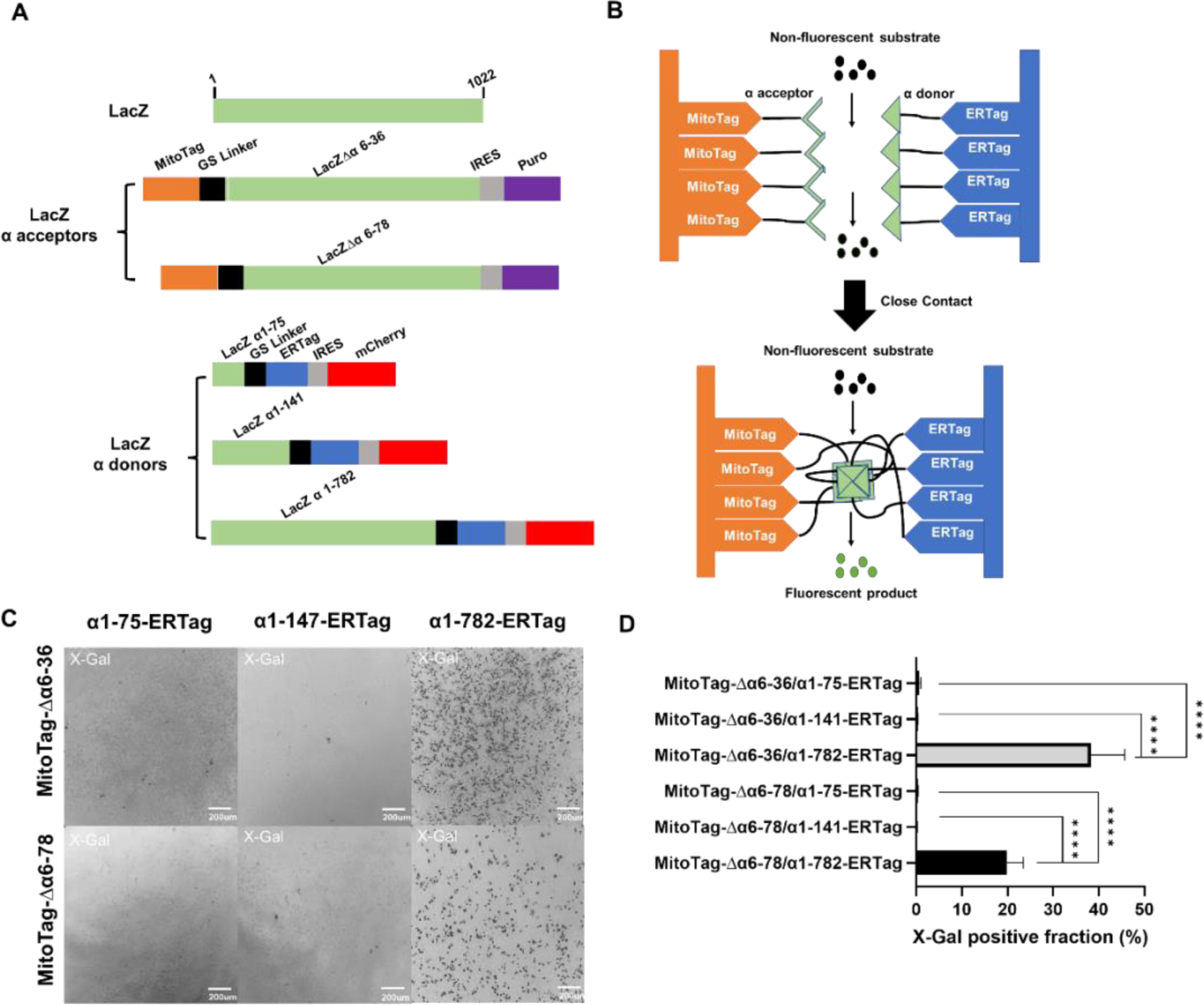
Constructs and concept of split LacZ based mitochondria-ER contact site (MERCS) reporter. (**A**) Schematic of the two mitochondria-targeted LacZ α acceptors (Δα6-36, Δα6-78) and the three ER-targeted LacZ α donors (α1-75, α1-141, and α1-782). The top green rectangle indicates the full-length LacZ gene. MitoTag is derived from the N-terminal transmembrane sequence from Tomm70, and ERTag is derived from the C-terminal transmembrane sequence from UBE2J2. GS, glycine-serine; IRES, internal ribosomal entry sequence (**B**) Diagram of the SpLacZ-MERCS reporter concept. The LacZ fragments are targeted separately to the surfaces of the mitochondria and ER. At regions where mitochondria and ER form contact sites (bottom panel), the split LacZ fragments are brought close enough to allow formation of enzymatically active octamers. Non-fluorescent β-Gal substrate is then hydrolyzed into fluorescent products. (**C**) Phase contrast images of cells containing 6 pairs of split LacZ fragments after incubation with X-Gal. Images were taken with a 10X objective. (**D**) Quantification of (**C**). Particle analysis in ImageJ was used to measure the fraction of cells that were X-Gal positive. Data are shown as mean ± s.d. More than 50 cells were analyzed from each experiment; *n*=3. ****, *p*≤0.0001; ***, *p*≤0.001; **, *p*≤0.01; *, *p*≤0.05; ns, *p*≥0.05. Statistical analysis was performed with the Student’s *t*-test. See also Fig. S1.

With two mitochondria-targeted α acceptors and three ER-targeted α donors, six different pairwise combinations were tested in U2OS cells. In previous studies, these LacZ fragment pairs were shown to have low affinity for each other, and enzyme activity was reconstituted only when the protein fragments were fused to other proteins that physically interact^23^. Both the MitoTag-Δα6-36/α1-782-ERTag and α1-782-ERTag/Δα6-78-MitoTag pairs showed substantial complementation, as evidenced by X-Gal staining (Figure 1C, D and S1A). The MitoTag-Δα6-36/α1-782-ERTag pair showed the highest complementation and was used for further studies. Control cell lines expressing only individual LacZ fragments showed no ability to hydrolyze β-gal substrate (Figure S1B).

Using immunofluorescence on cells co-expressing LacZ MitoTag-Δα6-36 and LacZ α1-782-ERTag, we confirmed that LacZ MitoTag-Δα6-36 colocalized with the mitochondrial marker Tomm20, and LacZ α1-782-ERTag colocalized with the ER marker Calnexin (Figure 2A, B). We similarly evaluated the previously established split-GFP-based MERCS reporter system (referred to as SpGFP-MERCS hereafter), which uses the same TOMM70 and UBE2J2 targeting sequences to target GFP11 to mitochondria (Mitot–spGFP11) and GFP1-10 to the ER (spGFP1-10–ERt)^19^. In cells co-expressing Mitot–spGFP11 and spGFP1-10–ERt, the mitochondrially targeted fragment co-localized with Tomm20. However, the spGFP1-10–ERt fragment appeared in tubular structures that did not colocalize with Calnexin (Figure 2C, D). Instead, the spGFP1-10–ERt-positive tubules colocalized with Tomm20 (Figure 2E, F). The corresponding LacZ α1-782-ERTag fragment did not show this organellar mislocalization (Figure 3E, F). The mislocalization did not occur when spGFP1-10–ERt was expressed in the absence of MitoTag–SpGFP11 (Figure S2A, B), indicating that mislocalization was likely induced by the high binding affinity between GFP11 and GFP1-10^20,29^.

**Figure 2.**
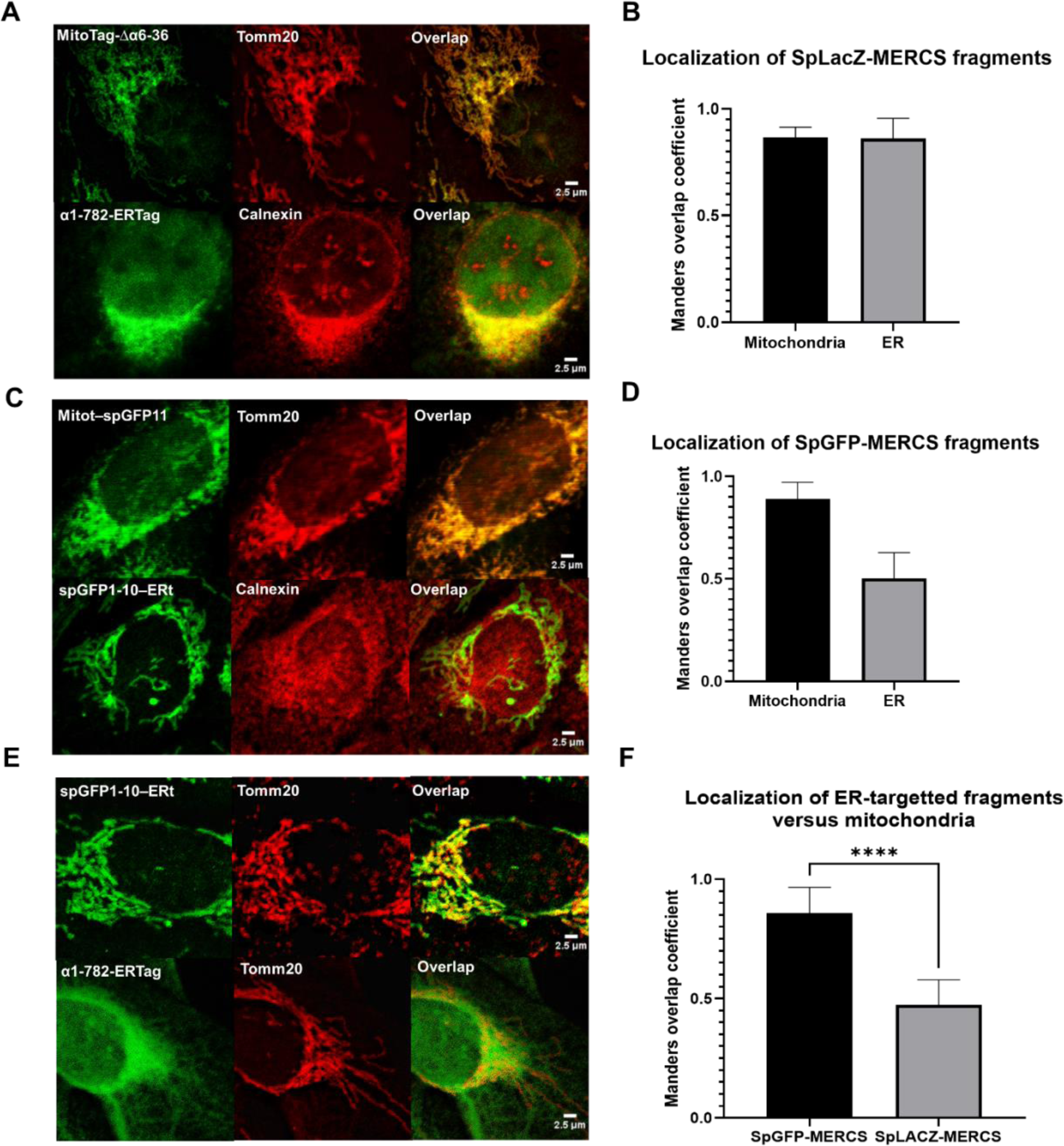
Comparison of the organellar targeting of SpLacZ-MERCS and SpGFP-MERCS fragments. (**A**) Targeting of SpLacZ-MERCS fragments in cells expressing both components of SpLacZ-MERCS. MitoTag-Δα6-36 (stained with Anti-LacZ) was compared with the mitochondrial marker Tomm20, and α1-782-ERTag (stained with Anti-V5) was compared with the ER marker Calnexin. (**B**) Manders coefficient analysis of experiment from (**A**). (**C**) Targeting of SpGFP-MERCS fragments in cells expressing both components of SpGFP-MERCS. MitoTag–SpGFP11 was compared with the mitochondrial marker Tomm20, and SpGFP1-10–ERTag was compared with the ER marker Calnexin. (**D**) Manders coefficient analysis of experiment from (**C**). (**E**) Comparison of the subcellular localizations of ER fragments from SpGFP-MERCS and SpLacZ-MERCS with the mitochondrial marker protein Tomm20. Both fragments were stained via the V5 protein tag. (**F**) Manders coefficient analysis of experiment from (**E**). In (**B**), (**D**), and (**F**), Manders overlap coefficient analysis is explained in the Methods. Data are shown as mean ± s.d. More than 20 cells were analyzed from each experiment; *n*=3. ****, *p*≤0.0001; ***, *p*≤0.001; **, *p*≤0.01; *, *p*≤0.05; ns, *p*≥0.05. Statistical analysis was performed with the Student’s *t*-test. See also Fig. S2.

**Figure 3.**
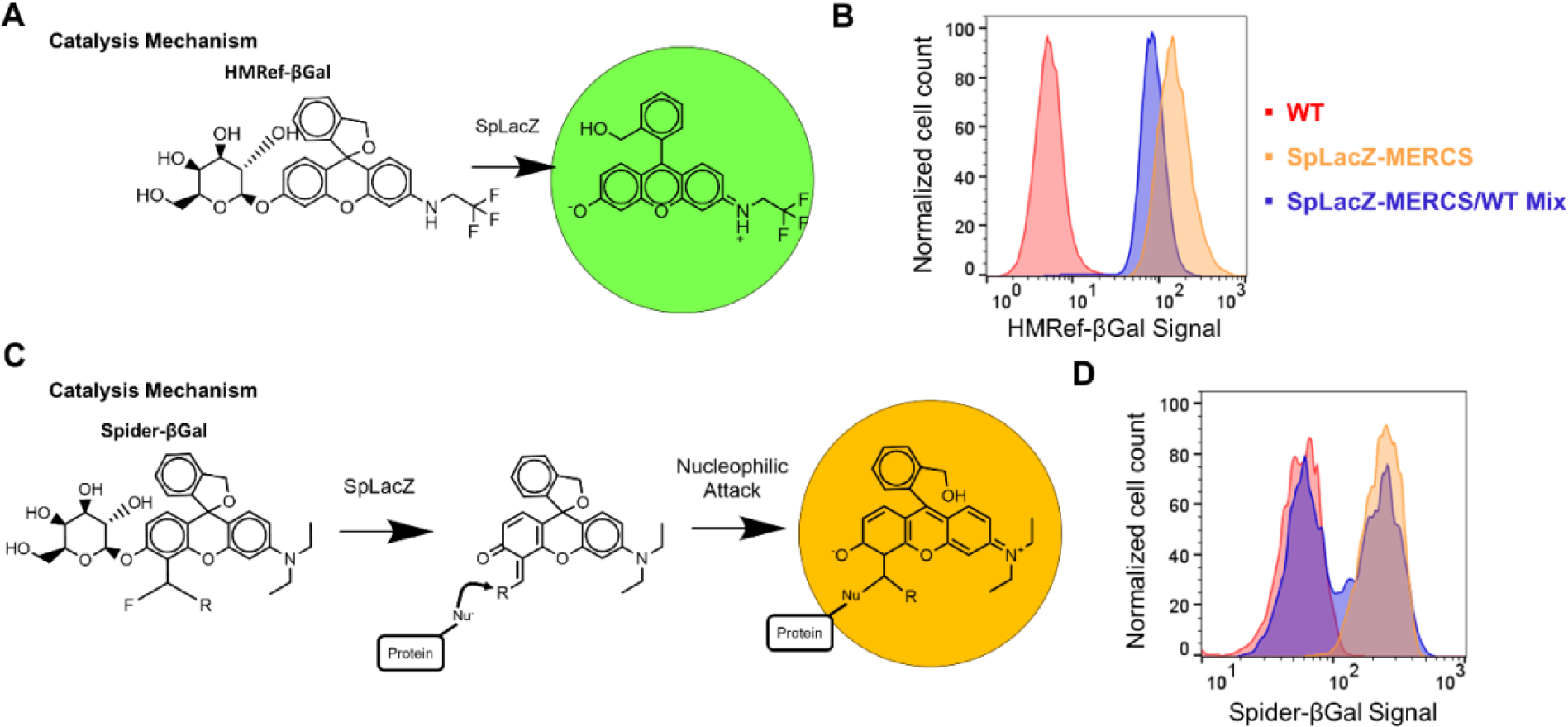
Spider-βGal is a preferred β-galactosidase substrate that enables fluorescence-based analysis of a mixed population. (**A**) Reaction mechanism for conversion of HMRef-βGal substrate into a soluble fluorescent product. The mechanism is reproduced from Asanuma et al^29^. Flow cytometry assay to determine cell autonomy of fluorescence signal. A mixed population (blue) of U2OS-WT and U2OS-SpLacZ-MERCS cells was analyzed by flow cytometry and compared to control and U2OS-SpLacZ-MERCS cells (1 μM HMRef-βGal, 4 h). The mixed population shows a single peak located between the control and U2OS-SpLacZ-MERCS cells. Reaction mechanism for conversion of Spider-βGal substrate into a reactive fluorescent product that covalently bonds with surrounding cellular proteins. The mechanism scheme is reproduced from Doura et al^30^. (**D**) Flow cytometry assay to determine cell autonomy of fluorescence signal. A mixed population (blue) of U2OS-WT and U2OS-SpLacZ-MERCS cells was analyzed by flow cytometry and compared to control and U2OS-SpLacZ-MERCS cells (0.25 μM Spider-βGal, 4 h). The mixed population shows two separate peaks, one aligned with U2OS-WT cells and one aligned with U2OS-SpLacZ-MERCS cells. For part (**B**) and (**D**), more than 20000 cells were analyzed for each sample; *n*=3. See also Fig. S3.

Because expression of artificial tethers can increase mitochondria-ER contact^30,31^, we tested whether the expression of the SpLacZ-MERCS reporter perturbed normal mitochondria-ER interactions. Live-cell imaging of cells expressing the SpLacZ-MERCS reporter showed no change in the levels of colocalization between the mitochondria and ER, as assessed by Tomm20 and Calnexin staining (Figure S2C, D).

### Spider-βGal substrate enables single-cell MERCS measurement

HMRef-βGal is a LacZ substrate that produces green fluorescence in live cells upon hydrolysis by β-galactosidase (structure and catalysis mechanism shown in Figure 3A)^32^. After incubation with HMRef-βGal, cells expressing SpLacZ-MERCS showed increased fluorescence compared to wild-type control cells. However, when a 1:1 mixture of the control cells and SpLacZ-MERCS-expressing cells was analyzed, only a single, intermediate peak appeared on flow cytometry (Figure 3B). The presence of the intermediate peak suggests that the fluorescent product leaks out of SpLacZ-MERCS-expressing cells and is taken up by control cells. This cell retention problem indicates that the HMRef-βGal substrate is not suitable for measuring the MERCS level in individual cells within a population.

To circumvent this problem, we tested Spider-βGal, an alternative β-galactosidase substrate whose cleavage product is a reactive quinone methide intermediate that reacts with cellular proteins and becomes immobilized^33^. Figure 3C shows the mechanism of action and structure of Spider-βGal. We confirmed that the SpLacZ-MERCS-expressing cells, in contrast to cells expressing a single SpLacZ fragment, converted Spider-βGal to its fluorescent state (Figure S3). Importantly, when we mixed an equal number of control cells with SpLacZ-MERCS cells, two distinct peaks were found that corresponded to the individual cell populations on flow cytometry (Figure 3D). These results suggest that Spider-βGal has no cell leakage and can be used for fluorescence-based analysis of SpLacZ-MERCS.

### The SpLacZ-MERCS reporter detects drug-induced MERCS defects

To test whether SpLacZ-MERCS can detect differences in MERCS levels caused by drugs, we examined the effect of oligomycin A, CCCP, and Mdivi-1 on mitochondria-ER colocalization and the SpLacZ-MERCS signal. Oligomycin A is an ATP synthase inhibitor that causes mitochondrial fission, a process that involves wrapping of the ER around mitochondrial tubules to cause constriction^34^. CCCP (carbonyl cyanide m-chlorophenyl hydrazone) disrupts the mitochondrial membrane potential and also induces mitochondria fission. Mdivi-1 (Mitochondrial division inhibitor 1) is an inhibitor of DRP1 with additional effects on Complex I^35,36^. As expected, oligomycin A and CCCP treatment resulted in substantial mitochondrial fragmentation associated with an increase in mitochondria-ER contacts, as measured by MitoTracker/ER-Tracker colocalization. In contrast, Mdivi-1 treatment resulted in a hyperfused mitochondria network with reduced mitochondria-ER colocalization (Figure 4A, B).

**Figure 4.**
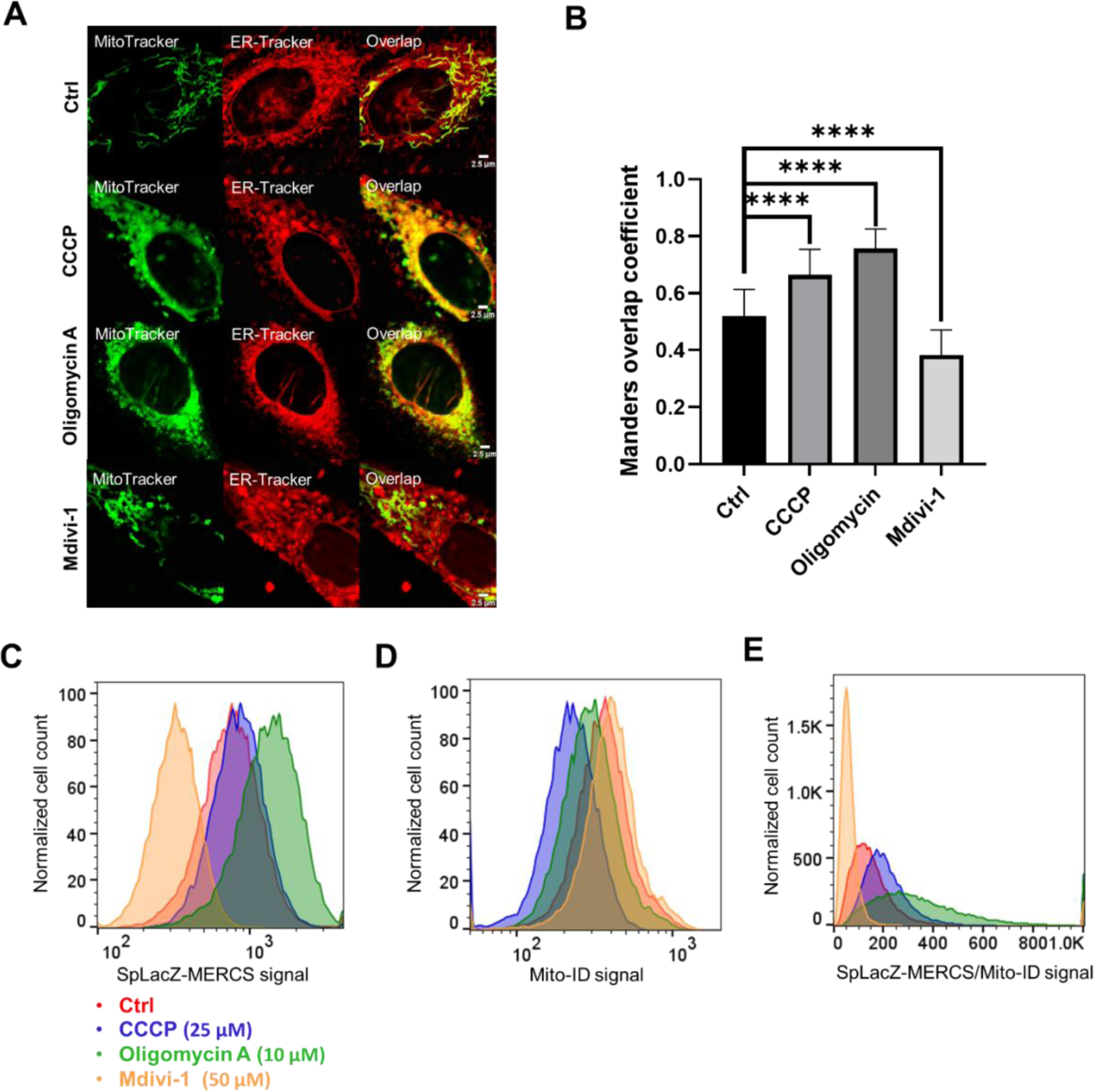
Effect of mitochondrial drugs on SpLacZ-MERCS signal. (**A**) Effect of mitochondrial drugs on colocalization of mitochondria and ER. The U2OS-SpLacZ-MERCS cell line was treated CCCP (25 μM, 4 h), oligomycin A (10 μM, 4 h), Mdivi-1 (50 μM, 4 h), or vehicle. Mitochondrial and ER structures were analyzed by staining with MitoTracker and ER-Tracker. (**B**) Quantification of ER/mitochondrial colocalization under CCCP treatment, oligomycin A treatment, Mdivi-1 treatment, and non-treatment conditions. Manders overlap coefficient analysis is explained in the Methods. Data are shown as mean ± s.d. More than 20 cells were analyzed from each experiment; *n* = 3. (**C**) Effect of selected drugs on the SpLacZ-MERCS signal. Flow cytometry was used to quantify the SpLacZ-MERCS reporter signal (0.25 μM Spider-βGal, 4 h). (**D**) Effect of mitochondrial drugs on the Mito-ID signal. Mito-ID stains mitochondria regardless of membrane potential and can be used as a proxy for mitochondrial mass. Flow cytometry was used to quantify Mito-ID signal from untreated cells and cells treated with CCCP, oligomycin A, or Mdivi-1. (**E**) Effect of mitochondrial drugs on the SpLacZ-MERCS signal after correction for mitochondrial mass. Cells were incubated with Spider-βGal and Mito-ID and analyzed by flow cytometry. The plot shows the Spider-βGal/Mito-ID ratio on the x-axis. For part (**C**-**E**), more than 20000 cells were analyzed for each sample; *n*=3. See also Fig. S4.

The SpLacZ-MERCS signal was dramatically increased by oligomycin A, moderately increased by CCCP, and decreased by Mdivi-1 (Figure 4C). CCCP is known to induce mitophagy and reduce mitochondrial content^37^. To normalize for mitochondrial content, we stained mitochondria with Mito-ID, a dye that marks mitochondria irrespective of membrane potential. There was a substantial reduction in Mito-ID staining after CCCP treatment (Figure 4D). Upon normalization for mitochondrial content, both CCCP and oligomycin A treatment caused substantial increases in the SpLacZ-MERCS signal, whereas Mdivi-1 caused a decrease (Figure 4E). Control experiments showed that these drugs did not interfere with the β-galactosidase hydrolysis reaction (Figure S4A). In contrast to our SpLacZ-MERCS, the SpGFP-MERCS was static, failing to show a response to any of the drug treatments (Figure S4B). We also found that the Spider-βGal incubation time or the Spider-βGal substrate level could be optimized to improve the signal-to-noise separation (Figure S4C, D).

### SpLacZ-MERCS signal is increased by overexpression of endogenous and artificial mitochondria-ER tethers

We tested whether our reporter responded to overexpression of native mitochondria-ER tethers. Three of the most well-characterized mitochondria-ER tethers are VAPB, PTPIP51, and PDZD8. VAPB and PTPIP51 are interacting proteins that localize to the ER and mitochondria, respectively, and are known to facilitate calcium transfer, lipid transfer, and regulation of mitochondria quality control^10,11,13,15^. PDZD8 is an integral ER membrane protein that has also been found to be important for mitochondria-ER contacts^12^. Overexpression of either VAPB, PTIPIP51, or PDZD8 increased the activity of the SpLacZ-MERCS reporter. Cells overexpressing both VAPB and PTPIP51 showed an even greater increase in the SpLacZ-MERCS signal (Figure 5A and S5A).

**Figure 5.**
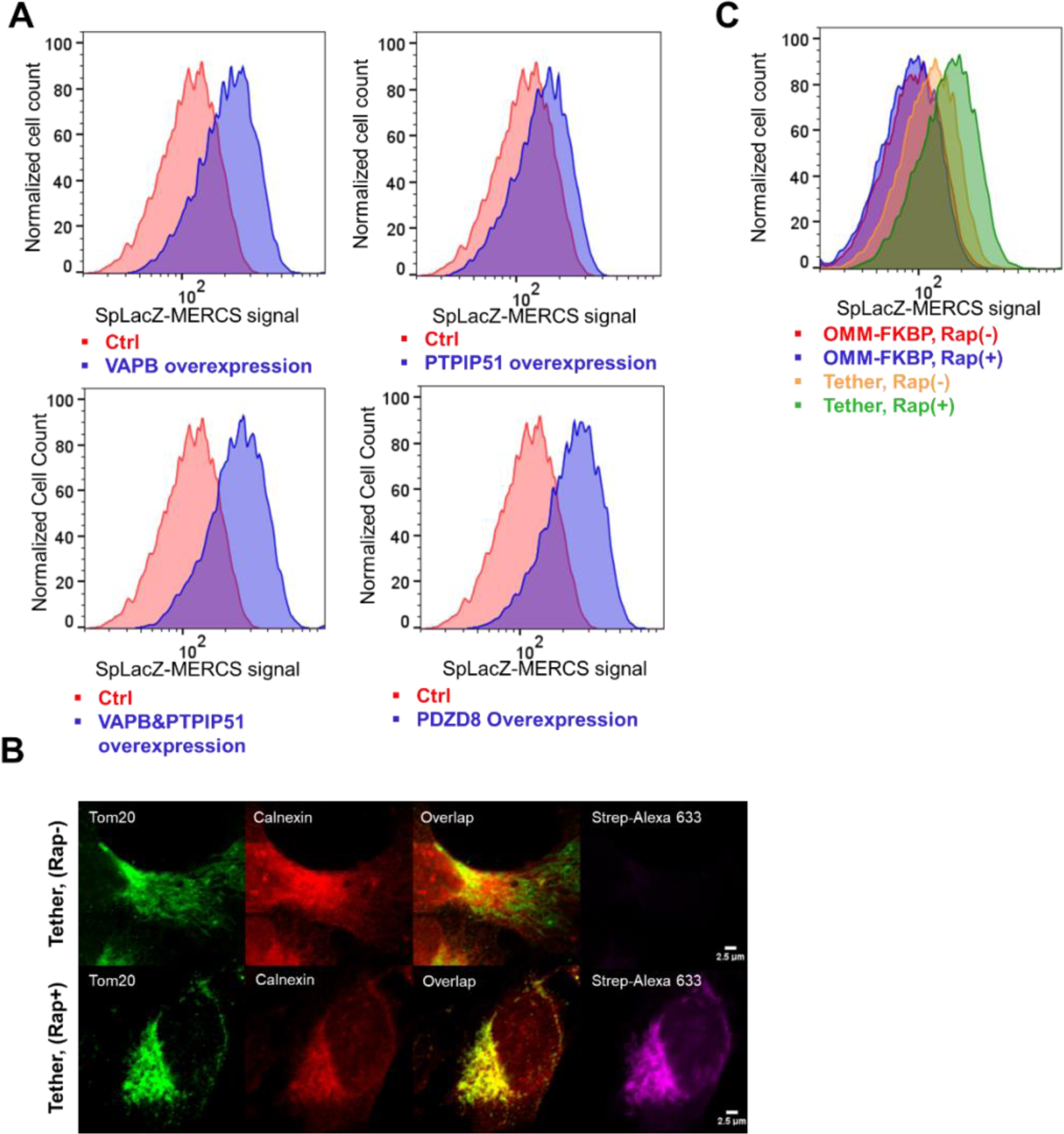
Effect of endogenous and artificial tethers on the SpLacZ-MERCS signal. (**A**) Effect of over-expressed endogenous tethers on the SpLacZ-MERCS signal. Flow cytometry was used to quantify the SpLacZ-MERCS signal (0.25 μM Spider-βGal, 4 h) upon stable expression of VAPB, PTPIP51, both VAPB and PTPIP51, and PDZD8. (**B**) Effect of artificial tether on mitochondria/ER colocalization. Cells expressed the split-TurboID FKBP-FRB system (tether). Upon rapalog (Rap) addition (bottom panel), mitochondria and ER are artificially tethered. ER and mitochondria colocalization were analyzed by immunofluorescence against Calnexin and Tomm20, respectively. Rapalog treatment also reconstitutes biotinylation activity^37^, which was detected by staining with streptavidin-Alexa Fluor 633. (**C**) Effect of artificial tethering on the SpLacZ-MERCS signal. Flow cytometry was used to quantify the SpLacZ-MERCS reporter signal (0.25 μM Spider-βGal, 4 h) in cells expressing the split-TurboID FKBP-FRB system (Tether) or only one component as a control (OMM-FKBP). Rapalog addition was used to mediate tethering in the former. For part (**A**) and (**C**), more than 20000 cells were analyzed for each sample; *n*=3. See also Fig. S5.

We then investigated the effect of artificial tethering of mitochondria-ER membranes on the SpLacZ-MERCS signal. The split-TurboID FKBP-FRB system^38^ is an artificial tether in which one component (OMM-FKBP) is localized to the mitochondrial outer membrane, and the other component (ER-FRB) is localized to the ER membrane. Strong association of the two components is triggered by rapalog, a small molecule that mediates binding between FKBP and FRB (Figure S5B). Rapalog also results in the reconstitution of TurboID, an engineered biotin ligase^38^. Upon addition of rapalog to cells expressing split-TurboID FKBP-FRB and SpLacZ-MERCS, we observed an increase in mitochondria-ER colocalization, biotinylation activity, and the SpLacZ-MERCS signal (Figure 5B, C).

### SpLacZ-MERCS signal is reduced by the knockdown of endogenous MERCS tethers

As a final test, we investigated the effect of knocking down native mitochondria-ER tethering factors. Using CRISPRi, we performed knockdowns of VAPB, PTPIP51 and PDZD8 in cells expressing either SpLacZ-MERCS or SpGFP-MERCS. For each tethering protein, successful knockdown for two gRNAs was confirmed by Western blotting (Figure S6A). Knockdown of VAPB, PTPIP51, or PDZD8 all resulted in lower SpLacZ-MERCS signal by flow cytometry (Figure 6A). However, the same knockdowns did not affect the SpGFP-MERCS signal (Figure 6B). For each of these knockdowns, mitochondrial mass, as measured by Mito-ID analysis, was not affected (Figure S6B).

**Figure 6.**
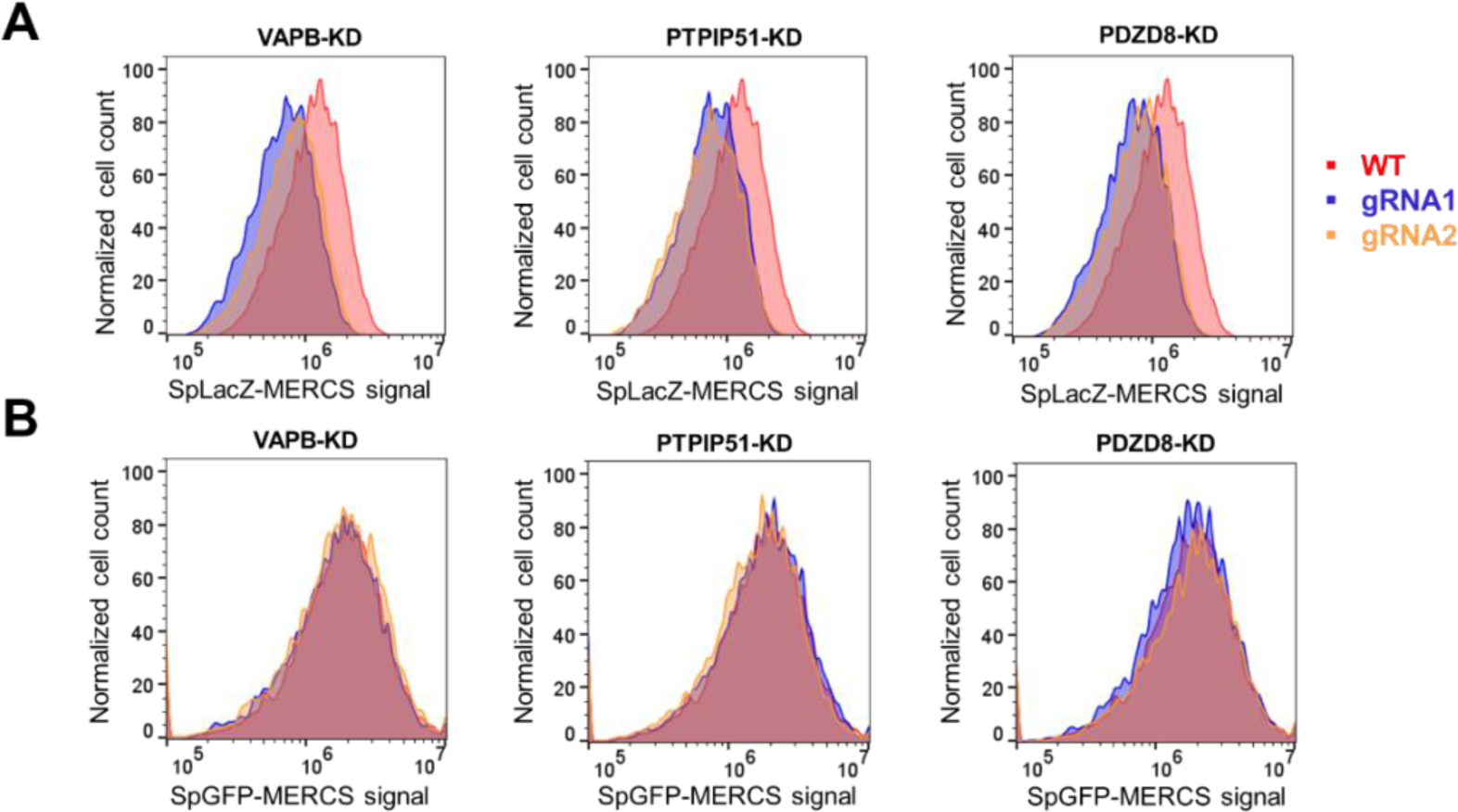
Effect of disruption of endogenous tethers on the SpLacZ-MERCS signal. (**A**) Effects of VAPB, PTPIP51, and PDZD8 knockdowns on the SpLacZ-MERCS signal. Flow cytometry was used to quantify the SpLacZ-MERCS signal (0.25 μM Spider-βGal, 4 h). (**B**) Effects of tether knockdowns on the SpGFP-MERCS reporter signal. Flow cytometry was used to quantify SpGFP-MERCS signal. For part (**A**) and (**B**), more than 20000 cells were analyzed from each sample; *n*=3. See also Fig. S6.

### SpLacZ-MERCS can accurately track MERCS dynamics

To test the ability of SpLacZ-MERCS to monitor MERCS dynamics, we performed time-lapse confocal imaging to evaluate the dynamics of the SpLacZ-MERCS signal against that of ER-mitochondria colocalization. Representative early-stage images of SpLacZ-MERCS show puncta formation at the interface between mitochondria and ER (Figure 7A). Untreated cells show fluctuations in mitochondria-ER colocalization that correlated well with the dynamics of the SpLacZ-MERCS signal (Figure 7B, D). Upon treatment of cells with oligomycin A to induce mitochondrial fission, cells showed a time-dependent increase in mitochondria-ER colocalization that correlated well with the increase in the SpLacZ-MERCS signal (Figure 7C, D). Under both conditions, the dynamics of the SpLacZ-MERCS signal and mitochondria/ER colocalization were highly correlated. (Figure 7D). The range of the SpLacZ-MERCS signal dynamics has a magnitude similar to that of the mitochondria-ER colocalization under both conditions (Figure 7E, F). Thus, our data suggests that SpLacZ-MERCS is capable of accurately capturing MERCS dynamics.

**Figure 7.**
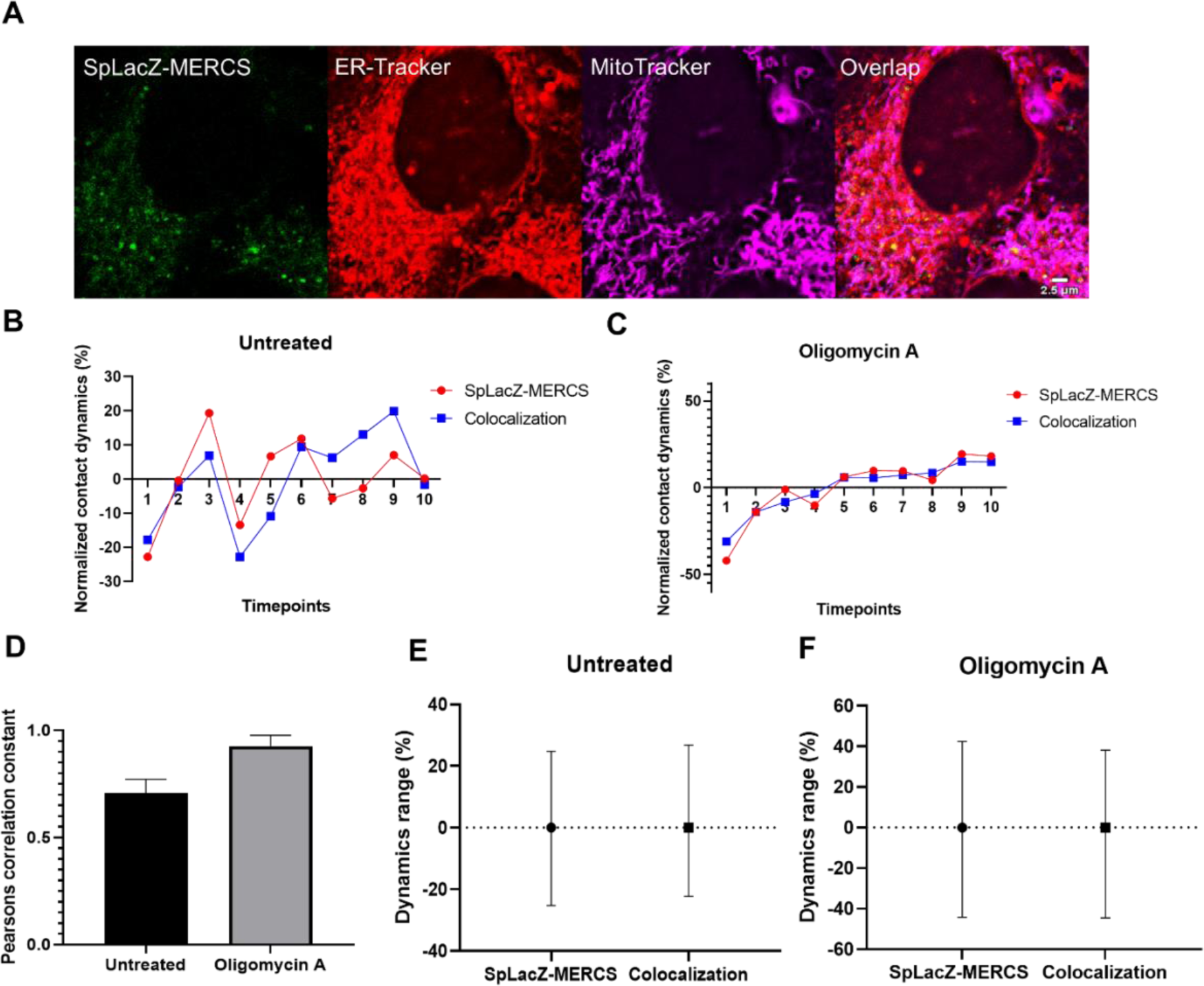
Tracking of MERCS dynamics by SpLacZ-MERCS reporter. (**A**) Representative still frames from a time-lapse movie of cells harboring the SpLacZ-MERCS reporter and treated with Spider-βGal (0.25 μM). (**B**) Comparison of contact dynamics measured by the SpLacZ-MERCS reporter versus mitochondria/ER colocalization in untreated cells. For the SpLacZ-MERCS reporter, "normalized contacts dynamics" is calculated at each timepoint as the difference in fluorescence intensity between the last and the current timepoint, normalized to the mean fluorescence value (Eq.1, 3). For mitochondria-ER colocalization, "normalized contacts dynamics" at each timepoint is the average of the Manders coefficient for mitochondria-ER colocalization for the last and current timepoint, normalized to the mean value (Eq.2, 4). Negative values indicate decreasing MERCS levels; zero indicates unchanged MERCS levels; positive values indicate increasing MERCS levels. The plots indicate the fluctuations in MERCS dynamics occurring during normal culture. (**C**) Comparison of contacts dynamics measured by the SpLacZ-MERCS reporter versus mitochondria/ER colocalization, after oligomycin A addition. Note that both measurements show progressive increases in MERCS content. (**D**) Correlation of the dynamics of the SpLacZ-MERCS signal to that of mitochondria/ER colocalization. Pearson correlation constants are shown for untreated cells and cells treated with oligomycin A. Data are shown as mean ± s.d. (**E**) The ranges of the contact dynamics measured by SpLacZ-MERCS and mitochondria/ER colocalization in untreated cells. (**F**) The ranges of the contact dynamics measured by SpLacZ-MERCS and mitochondria/ER colocalization after oligomycin A addition. For **(B-F)**, the Manders overlap coefficients analysis is explained in the Methods. For (**D-F**), more than 5 cells were analyzed from each experiment; *n* = 3.

## Conclusions

MERCS are dynamic inter-organellar interfaces that coordinate the activities of the ER and mitochondria, including calcium homeostasis, lipid biosynthesis, mitochondrial dynamics, and mitochondrial quality control. Considering the importance of this topic, it is critical to have an accurate MERCS reporter that overcomes the limitations of current systems. In this study, we developed the first MERCS reporter system that generates a fluorescent signal that accumulates over time and, therefore, integrates the history of the MERCS signal. SpLacZ-MERCS combined with the Spider-β Gal substrate enabled the accurate analysis of MERCS levels within individual cells of a population. We identified a LacZ α acceptor and α donor pair that functions well together when targeted to mitochondria and ER respectively. Each reporter fragment was cleanly trafficked to its respective compartment. Although correct organellar targeting may seem trivial, we found that the ER component of the SpGFP-MERCS reporter has a high degree of mislocalization to mitochondria. This mislocalization is likely caused by the essentially irreversible binding of split-GFP fragments.

The SpLacZ-MERCS reporter responded well to drug and genetic manipulations that affect the interaction between the mitochondria and ER. The drugs CCCP, oligomycin A, and Mdivi-1 all affected mitochondria-ER colocalization and have a corresponding effect on the SpLacZ-MERCS signal. Overexpression of the well-characterized tethers VAPB, PTPIP51, and PDZD8 each resulted in an increase in the SpLacZ-MERCS signal. Expression of an artificial FKBP/FRB-based mitochondria-ER tethering system also resulted in a significant increase in SpLacZ-MERCS signal upon chemically induced dimerization. Knockdowns of VAPB, PTPIP51, or PDZD8 reduced the SpLacZ-MERCS signal. In contrast, the SpGFP-MERCS reporter failed to respond to the knockdown of known tethers. Time-lapse studies showed that the SpLacZ-MERCS reporter is able to track dynamic changes in the MERCS level, whereas the signal of the SpGFP-MERCS reporter is static.

The integrative signal produced from the SpLacZ-MERCS reporter is advantageous in evaluating individual cells for their overall MERCS levels. Mitochondria-ER interactions are dynamic, and cells are expected to have fluctuations in their MERCS level due to factors such as organelle motility, organelle shaping, and cell cycle. These fluctuations complicate the ability of other MERCS reporters to score cells as having high or low levels of MERCS, whereas SpLacZ-MERCS can accurately reflect the overall MERCS level. This system enables new opportunities for the high-throughput screening of genes or drugs that regulate MERCS levels. A caveat is that our MERCS reporter cumulatively records the MERCS level over time, which prevents the visualization of exact contact sites. Our methodology may be applicable to studying the crosstalk of other organelles, like mitochondria-peroxisome and mitochondria-lysosome interactions.

## Methods

### Antibodies and reagents

Primary antibodies: Tomm20 (Santa Cruz BioTech, sc-17764), Calnexin (Proteintech, 66903-1-Ig), Myc (Sigma, C3956), FLAG M2 (Sigma, F1804-200UG), HA.11 (Covance, MMS-101R), PDZD8 (Proteintech, 25512-1-AP), VAPB (Proteintech, 14477-1-AP), PTPIP51 (Proteintech, 20641-1-AP).

Secondary antibodies: Goat anti-mouse IgG (H+L)-HRP (Jackson ImmunoResearch, 115-035-003), Goat anti-rabbit IgG (H+L)-HRP (Jackson ImmunoResearch, 111-035-003), Donkey anti-mouse IgG AlexaFluor 405 (abcam, ab175658), Donkey anti-mouse IgG AlexaFluor 488 (Invitrogen, A21202), Donkey anti-rabbit IgG AlexaFluor 555 (Invitrogen, A32794), Goat anti-rabbit IgG AlexaFluor 633 (Invitrogen, A21070).

Chemicals: carbonyl cyanide 3-chlorophenylhydrazone (CCCP) (Sigma-Aldrich, C2759), Mdivi-1 (Sigma-Aldrich, M0199), oligomycin A (Sigma-Aldrich, O4876), Spider-βGal (Dojindo, SG02), rapalog (Takara Bio, 635056), BioTracker 519 Green β-Gal Dye (Millipore Sigma, SCT025)

### Plasmid construction

Primer sequences are listed in Supplemental File S1. For the construction of LacZ donors plasmids, LacZ α1-75, α1-147, and α1-782 were amplified using the common forward primer Comdon-F and the reverse primers S-R, M-R, and L-R. The ER targeting sequence was amplified from plx304-spGFP1-10-Ert^19^ with primers ER-F and ER-R. NotI/MfeI-digested LacZ donor fragment and MfeI/BamHI-digested ERTAG were ligated with NotI/BamHI-digested backbone (PQCXIP-mCherry retroviral vector).

For the construction of LacZ acceptors plasmids, LacZ Δα6-36, and Δα6-78 were amplified respectively using the forward primers 1-F and 2-F, and a common reverse primer Comrec-R. The mito targeting sequence was amplified from pLVX -Mitot-spGFP11×2^19^ with primers Mito_F and Mito_R. The NotI/MfeI-digested LacZ acceptor fragment and MfeI/NotI-digested MitoTag were ligated with NotI/BamHI-digested backbone (PQCXIP-PURO retroviral vector).

For the construction of VAPB, PTPIP51, and VAPB/PTPIP51 expressing plasmids. The pUltra (Addgene Plasmid #24129) lentiviral vector is used as the backbone. The selection marker was modified from GFP to hygromycin by amplifying the hygromycin resistance gene with HYG_F and HYG_R and ligating into AgeI/BsrGI-digested pUltra. VAPB was amplified with VA_F and VA_R; PTPIP51 was amplified with PTP_F and PTP_R. VAPB and PTPIP51 ORFs were inserted into pUltra_Hyg. For the construction of PDZD8 expressing plasmid, PDZD8-3XHA was amplified from pCAG-PDZD8HA with PDZ_F and PDZ_R and inserted into PQCXIP-Neo digested with NotI/AgeI.

The gRNA plasmids were constructed by inserting annealed oligos into the lentiviral CRISPRia-v2 backbone (Addgene, 84832) at the BstXI/BlpI sites. For two gRNAs targeting VAPB, YP.190 and YP.191, and YP.192 and YP.193 were respectively annealed. For the two gRNAs targeting PTPIP51, YP.196 and YP.197, and YP.198 and YP.199 were respectively annealed. For the two gRNAs targeting PDZD8, YP.202 and YP.203, and YP.204 and YP.205 were respectively annealed. The gRNA control had the following protospacer sequence: gctcggtcccgcgtcgtcgg.

### Cell culture and generation of stable cell lines

U2OS and HEK293T cells were grown in DMEM supplemented with 10% fetal bovine serum (FBS), 2 mM glutamine, and 1% penicillin-streptomycin, at 37°C with 5% CO2. To produce retrovirus, HEK293T cells were transfected by calcium phosphate with packaging plasmids (pVSV-G and pUMVC) and retroviral constructs. For lentivirus production, HEK293T cells were transfected with pVSV-G, pΔ8.9, and lentiviral constructs. Fresh media was added 12 h after transfection. 48 h after transfection, the supernatant was collected and passed through a 0.45um syringe filter to remove cell debris. HeLa cells or K562 cells were transduced in the presence of 8 μg/mL polybrene (Sigma, H9268). To select for transduced cells, puromycin (1 ug/mL) or hygromycin (80 µg/ml) was applied at least 3 days or 7 days, respectively.

### Flow cytometry

**F**low cytometry analysis was performed with the S3e™ Cell Sorter (488/561 nm). For experiments knocking down endogenous MERCS tethers, BFP positive cells were sorted on a CytoFLEX S (Beckman Coulter).

To prepare for flow cytometry, cells were trypsinized, neutralized with Fluorobrite complete medium, and spun down at 300 *g* for 8 mins. Cells are washed once with ice-cold Fluorobrite complete medium and resuspended in Fluorobrite complete medium containing 20 mM HEPES before analysis and sorting. All flow cytometry data were analyzed in FlowJo v10.8 Software (BD Life Sciences).

### Immunostaining and live cell Imaging

For immunofluorescence imaging, cells were fixed in 4% paraformaldehyde in PBS, washed three times with PBS, and permeabilized with 0.1% Triton X-100 for 30 mins. Cells were washed three times with PBS and blocked in PBS containing 10% FBS for 30 mins. Fixed cells were further incubated overnight in the cold room with primary antibodies. Cells were washed with PBS three times and incubated with secondary antibodies at room temperature for 1 h and washed four times with PBS before imaging.

For measurements of mitochondria-ER colocalization, cells were washed and incubated with complete DMEM medium containing 200 nM MitoTracker Deep Red FM and 500 nM ER-Tracker Blue-White DPX for 30 mins at 37°C. Cells were washed three times and incubated in complete Flourbrite medium containing 10% fetal bovine serum, 2 mM glutamine, and penicillin-streptomycin before live cell imaging.

For both Immunofluorescence imaging and live cell imaging, images were obtained with a Zeiss LSM 710 confocal microscope (Carl Zeiss). Images were analyzed using ImageJ.

### Manders overlap coefficient analysis

For Figure 3B, Manders overlap coefficients measure the fraction of Tomm20 signal that overlapped with MitoTag-Δα6-36 signal, and the fraction Calnexin signal that overlapped with α1-782-ERTag signal. For Figure 3D, Manders overlap coefficients measure the fraction of Tomm20 signal that overlapped with Mitot–spGFP11 signal, and the fraction of Calnexin signal that overlapped with spGFP1-10–ERt signal. For Figure 3F, Manders overlap coefficients measure the fraction of spGFP1-10–ERt signal that overlapped with Tomm20 signal, and the fraction of α1-782-ERTag signal that overlapped with Tomm20 signal.

For Figure 5B and Figure 8B-F, Manders overlap coefficients measure the fraction of ER-Tracker signal that overlapped with MitoTracker signal.

For Figure S2B, Manders overlap coefficients measured the fraction of Calnexin signal that overlapped with spGFP1-10–ERt signal. For Figure S2D, Manders overlap coefficients measured the fraction of Calnexin signal that overlapped with the Tomm20 signal.

### Drug Treatments

The following drug concentrations were used: CCCP, 10 μM; Mdivi-1, 50 μM; oligomycin A1, 10 μg/ml. Cells were incubated with these drugs for four hours before flow cytometry or live-cell imaging.

### Analysis of SpLacZ-MERCS dynamics

To analyze the dynamics of the SpLacZ-MERCS signal, eleven images from a time-series were used for each cell. *F*_*spLacZ*_ (*n*) is defined as cell fluorescence level at timepoint n measured by SpLacZ-MERCS. *D*_*spLacZ*_ (*n*) is the change in contact dynamics based on the SpLacZ-MERCS reporter signal at timepoint n. *μ*_*D*_*spLacZ*__ is defined as the mean value of the ten *D*_*spLacZ*_ datapoints. 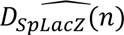 is defined as *D*_*spLacZ*_ at timepoint n normalized to the mean *D*_*spLacZ*_ value.

*M*(*n*) is defined as Manders overlap coefficient between mitochondria and ER at timepoint n. *D*_*Overlap*_(*n*) is the change in contact dynamics calculated based on mitochondria/ER colocalization at timepoint n. *μ*_*DOverlap*_ is defined as the mean value of the ten *D*_*Overlap*_ datapoints. 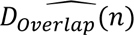 is defined as *D*_*Overlap*_ at timepoint n normalized to the mean *D*_*Overlap*_ value.

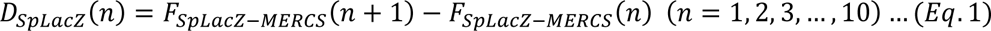

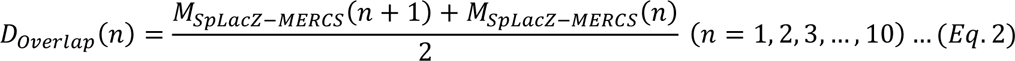

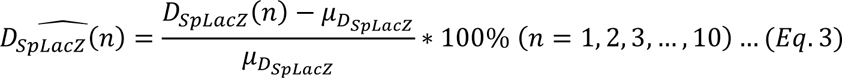

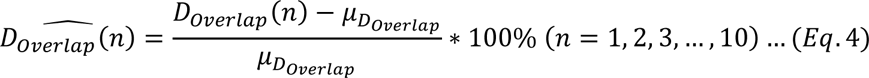

### Rapalog-induced mitochondria-ER tethering

The constructs pLX208 CMV sTurboID (C)-HA-FRB-ERM (Plasmid #153007) and pLX304 CMV OMM-FKBP-V5-sTurboID (N) (Plasmid #153006) were used to induce artificial tethering between mitochondria and ER. Cells were incubated with 500 nM rapalog and 50 μM of biotin for 24 h for immunofluorescence imaging. For flow cytometry cells were incubated with only rapalog.

### Statistical analysis

The statistical analysis was performed using GraphPad Prism 9. All data were shown as means±std. Statistical analysis among different groups was performed with the Student’s *t*-test. p≤0.0001; ***, p≤0.001; **, p≤0.01; *, p≤0.05; ns, p≥0.05.

## Supporting information

Supplemental figures: Supplemental figures S1-S6 are provided as a composite PDF file. Supplemental file 1: A list of DNA primers used in this study.

## Abbreviations

ATP: adenosine triphosphate
BRET: bioluminescence resonance energy transfer
CCCP: carbonyl cyanide m-chlorophenyl hydrazone
ER: endoplasmic reticulum
FRET: fluorescence resonance energy transfer
GFP: green fluorescent protein
MERCS: mitochondria-endoplasmic-reticulum contact sites

## Author information

Corresponding author:

David C. Chan

Division of Biology and Biological Engineering

California Institute of Technology

1200 East California Blvd, MC114-96

Pasadena, CA 91125

https://orcid.org/0000-0002-0191-2154

Author:

Zheng Yang

Division of Biology and Biological Engineering

California Institute of Technology

1200 East California Blvd, MC114-96

Pasadena, CA 91125

https://orcid.org/0000-0001-8131-0868

## Author Contributions

ZY and DCC formulated the research plan, and ZY performed the experiments. ZY and DCC wrote the manuscript.

## Acknowledgement

This work was supported by NIH grant R35GM127147.

