## Supplemental Figures for "Development of a signal-integrating reporter to monitor mitochondria-ER contacts"

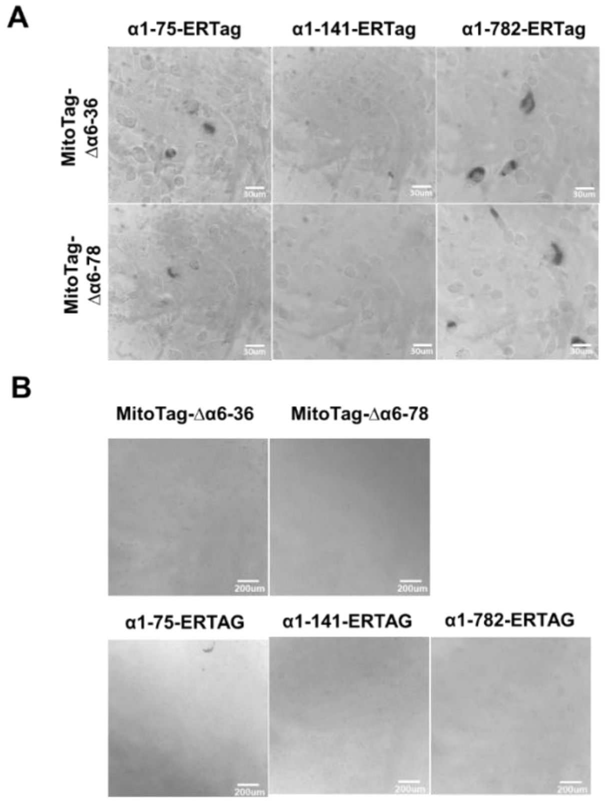


**Figure S1. Screening of LacZ fragment pairs targeted to mitochondria and ER.** (**A**) LacZ activity in cells containing split-LacZ reporters. Cells expressing the indicated split LacZ pairs were fixed, stained with X-Gal, and imaged with a 63X objective. (**B**) LacZ activity in cells expressing only one LacZ fragment. None of them produce a strong X-Gal signal. Images were taken with a 10X phase contrast objective.

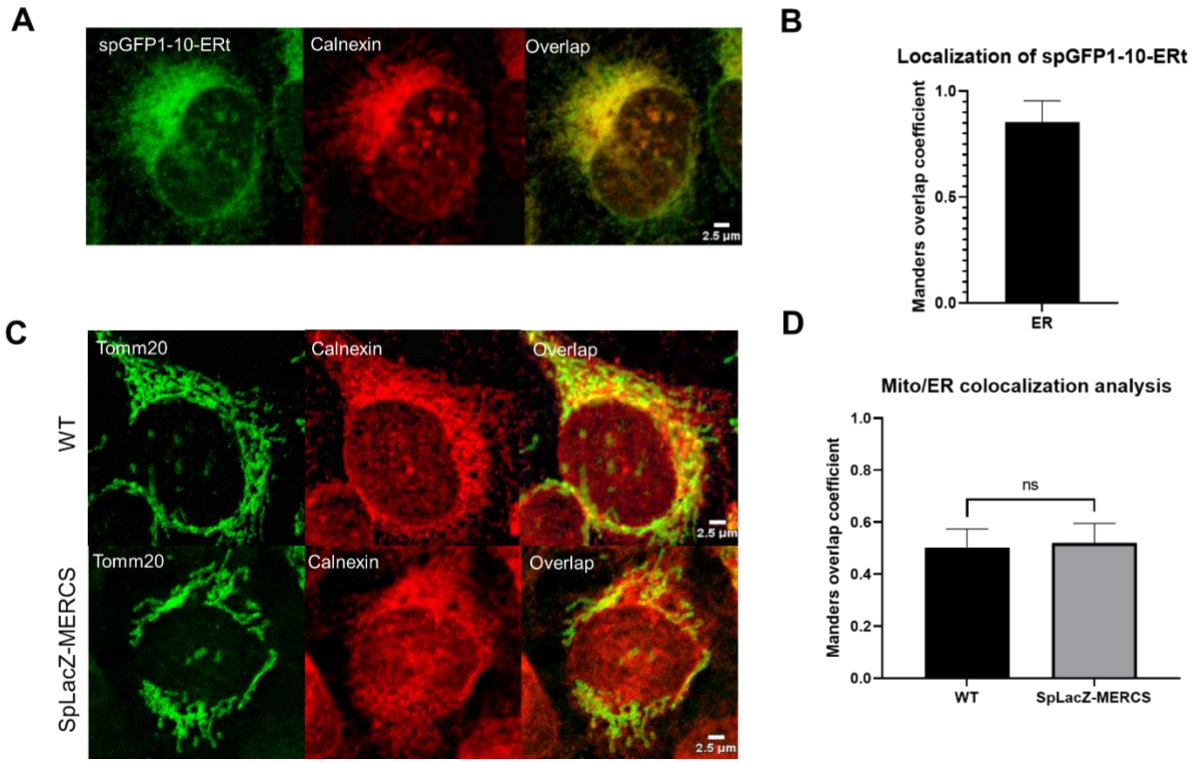


**Figure S2. spGFP1-10–ERt localization and Effect of SpLacZ-MERCS reporter on ER/mitochondria contacts.** (**A**) Targeting of spGFP1-10–ERt in cells when expressed alone. spGFP1-10–ERt was compared with the ER marker Calnexin. (**B**) Manders coefficient analysis of experiment from (**A**). (**C**) Colocalization between mitochondria and ER for wildtype U2OS cells and U2OS cell expressing the SpLacZ-MERCS reporter. Immunofluorescence was used to identify mitochondria (Tomm20) and ER (Calnexin). (**D**) Quantification of the colocalization between mitochondria and ER. Data are shown as mean ± s.d. More than 20 cells were analyzed from each experiment; $n$=3. The following $p$-value designations are used in all figures: ****, $p \le 0.0001$; ***, $p \le 0.001$; **, $p \le 0.01$; *, $p \le 0.05$; ns, $p \ge 0.05$. Statistical analysis was performed with the Student's $t$-test.

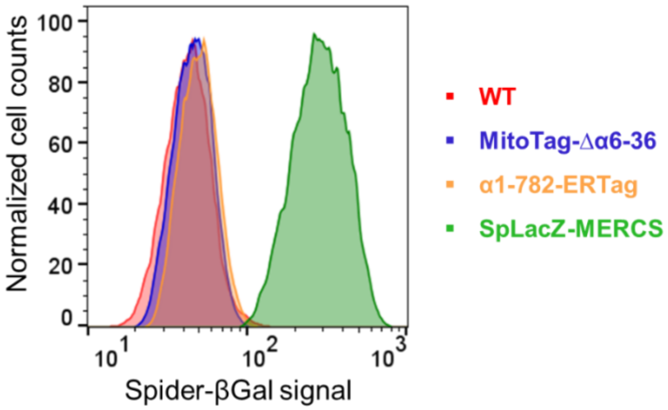


**Figure S3. Spider-βGal as a preferred β-galactosidase substrate for fluorescence-based analysis .** Spider-βGal fluorescence was analyzed by flow cytometry (0.25 μM Spider-βGal, 4 h). Only cells that harbor the full SpLacZ-MERCS reporter show a substantial fluorescent signal increase. More than 20000 cells were analyzed for each sample; *n*=3.

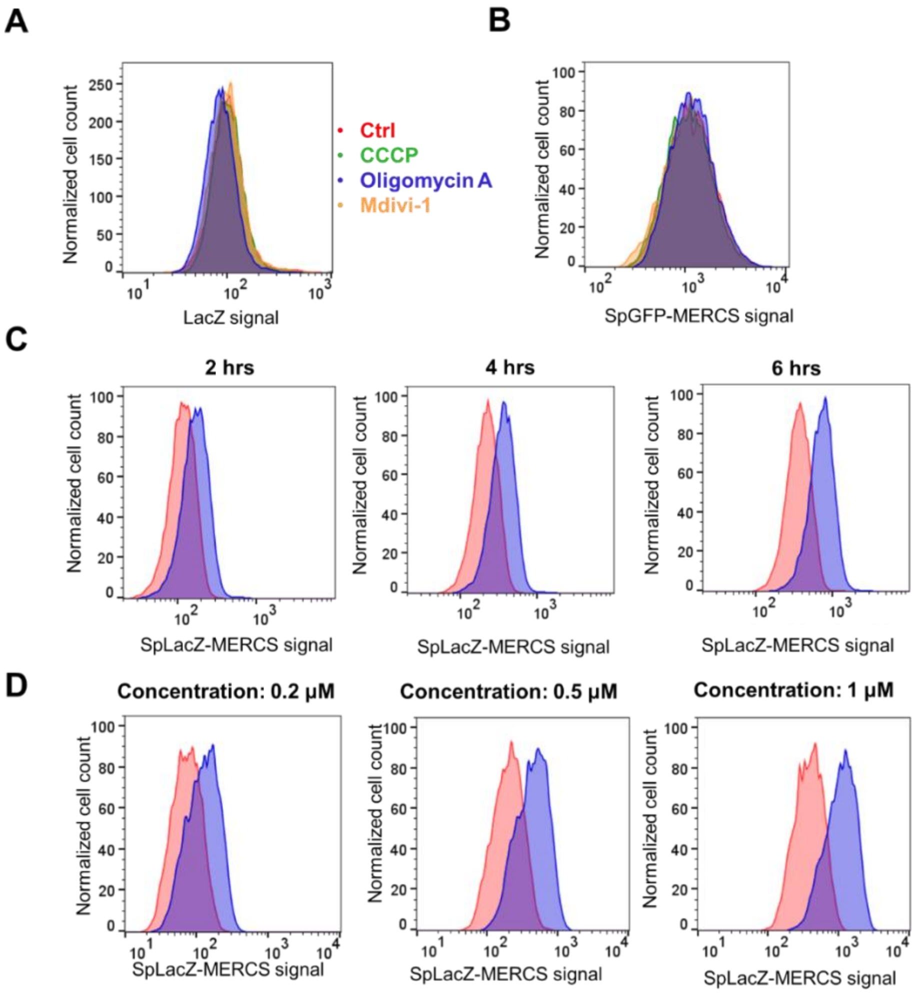


**Figure S4. Effect of mitochondrial drugs on the SpLacZ-MERCS reporter.** (**A**) Effect of mitochondrial drugs on cytosolic, full-length LacZ. Flow cytometry was used to quantify LacZ signal in cells treated with the indicated drug treatments (0.25 μM Spider-βGal, 4 h). (**B**) Effect of mitochondrial drugs on the SpGFP-MERCS reporter. Flow cytometry was used to quantify SpGFP-MERCS signal in cells treated with the indicated drug treatments. No significant differences in SpGFP-MERCS signal were observed between CCCP, oligomycin A, Mdivi-1 and control. (**C**) Effect of substrate incubation time on SpLacZ-MERCS signal. Flow cytometry was used to quantify the SpLacZ-MERCS signal (0.25 μM Spider-βGal) from cells with and without oligomycin A at different incubation times (2h, 4h and 6h). (**D**) Effect of substrate concentration on SpLacZ-MERCS signal (0.2, 0.5, 1.0 μM Spider-βGal). Cells with and without oligomycin A treatment were incubated with the indicated substrate concentrations. For part (**A**-**D**), more than 20,000 cells were analyzed from each sample; *n*=3.

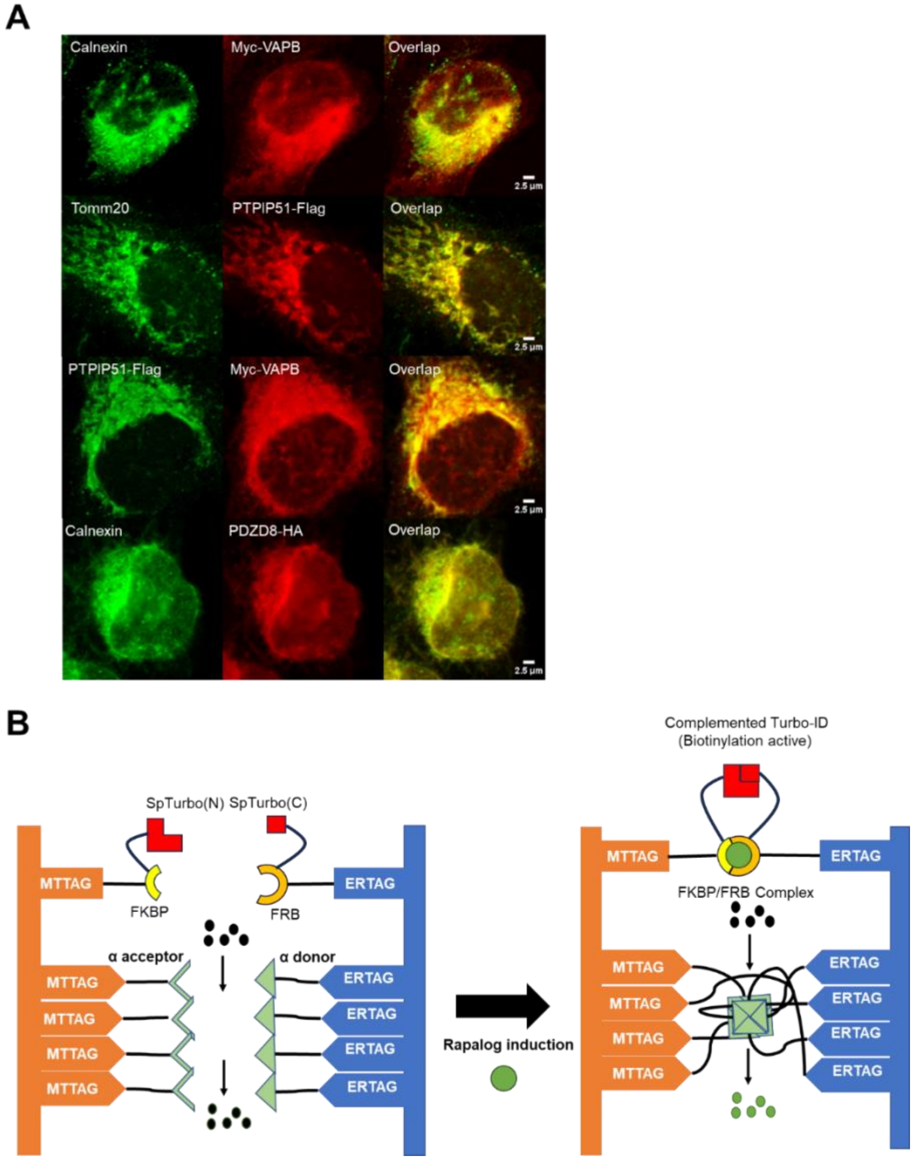


**Figure S5. Subcellular localization of over-expressed endogenous tethers and artificial tethering mechanism (A)** Localization of expressed VAPB, PTPIP51, and PDZD8. Images in third row show cells with co-expression of Myc-tagged VAPB and Flag tagged PTPIP51. **(B)** Diagram of the artificial induction of mitochondria-ER contact using split-TurboID FKBP-FRB constructs[37]. When rapalog is added, FKBP and FRB strongly interact to induce mitochondria-ER tethering, thereby activating the SpLacZ-MERCS system.

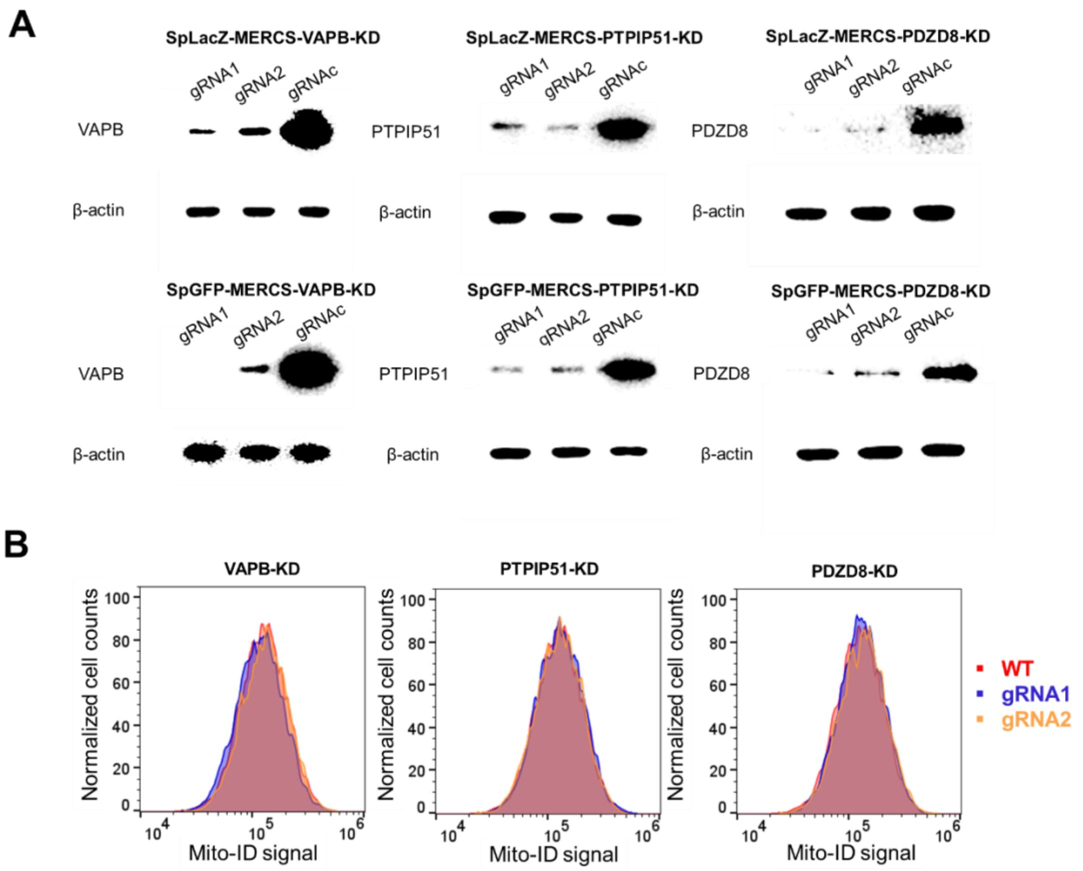


**Figure S6. Effect of knockdown of endogenous mitochondria-ER tethers.** (**A**) Western blot analysis of CRISPRi knockdown. Western blot was used to measure knockdown efficiency against VAPB, PTPIP51, and PDZD8. gRNAc is a non-targeting control gRNA. (**B**) Cells containing the indicated endogenous tether knockdown were stained with Mito-ID and analyzed by flow cytometry. More than 20000 cells were analyzed for each sample; *n*=3.
